# Multi-gene phylogeny and morphology of *Pleurotus* in Aotearoa New Zealand reveal a new variety of *Pleurotus pulmonarius*

**DOI:** 10.1101/2025.04.29.651148

**Authors:** David Hera, Jerry Cooper, Peter K. Buchanan, Manpreet K. Dhami, Ian A. Dickie

## Abstract

Increased demand of cultivated oyster mushrooms (*Pleurotus* species) in Aotearoa New Zealand has led to the importation of exotic species, which pose potential invasion risks. Gaps in taxonomic knowledge of this genus have complicated biosecurity decisions and cultivation efforts. To address this, we collected 84 wild and cultivated New Zealand *Pleurotus* specimens for multi-locus phylogenetic analysis (ITS, LSU, RPB1, RPB2, and Tef) and morphological examination. We describe *P. pulmonarius* var*. aotearoa* as a new variety indigenous to New Zealand, distinct from imported *P. pulmonarius* sens. str. We establish that *P. purpureo-olivaceus* has no anamorphic stage and falls outside the subgenus *Coremiopleurotus*, unlike *P. australis,* a species sometimes found on living trees. The close monophyletic relationship between *P. parsonsiae* and *P. djamor* underscores the need to reconsider the presence of the exotic *P. djamor* in the country. The refined species boundaries between the indigenous *P. australis, P. parsonsiae, P. pulmonarius* var*. aotearoa* and *P. purpureo-olivaceus* have important implications for conservation and biosecurity, and support the potential of using indigenous strains for cultivation in New Zealand.

## Introduction

Species importation for food and fodder can serve as important pathways for the inadvertent introduction and spread of invasive species (Bruce, 2018; Forseth & Innis, 2004; Shackleton et al., 2014). Island nations, such as Aotearoa New Zealand, are especially vulnerable to biological invasions, and thus manage this risk through strong biosecurity protocols by greatly restricting new species introductions, but biosecurity restrictions rely on clear and correct species definitions. The growing demand for cultivated mushrooms has resulted in movement of some strains internationally, at least one of which is now a major invasive species in North America (Veerabahu et al., 2024). An alternative approach to relying on new species introductions has been an increasing interest in cultivation of indigenous species, which are currently underutilised in local mushroom cultivation, and which could add value by leveraging the consumer appeal for indigenous foods. To adopt indigenous *Pleurotus* species for cultivation, the current gaps in taxonomic knowledge of this genus in New Zealand need to be addressed, particularly the relationship between local populations of species that have been historically identified as species originally described from the northern hemisphere (Bickford et al., 2007; Humphreys & Linder, 2009). This study aims to clarify the molecular phylogeny of indigenous and exotic *Pleurotus* species in New Zealand, addressing gaps in existing taxonomic descriptions and thereby informing local cultivation practices and biosecurity decisions.

*Pleurotus* (Pleurotaceae, Agaricales) is a genus of white rot wood decay fungi (Stamets, 2000) with more than 170 species names currently described globally on Index Fungorum (www.indexfungorum.org, accessed 11/02/2025). *P. ostreatus* (Jacq. ex Fr.) P.Kumm*., P. cornucopiae* (Paulet) Quél., *P. eryngii* (DC.) Quél., *P. djamor* (Rumph. ex Fr.) Boedijn and *P. pulmonarius* (Fr.) Quél. are some of the species commonly cultivated for food, fodder and bioactives (Cohen et al., 2002; Li et al., 2017; Royse et al., 2017). Oyster mushrooms are among the top three types of mushrooms by mass produced worldwide (Mortimer et al., 2021), along with shiitake and wood ear mushrooms.

Several indigenous species of *Pleurotus* are known in New Zealand, some of which are cultivated, alongside the introduced *P. djamor* and *P. pulmonarius*. Segedin et al. (1995) described six naturally occurring species. The authors’ species concept was largely based on the morphology, culture characteristics, and interfertility tests of four of the six species described. Historically, applying the biological species concept of reproductive isolation to fungi has been challenging. For example, some fungi exclusively reproduce asexually (Chethana et al., 2021), and other fungi are unculturable, making intercompatibility studies impossible (Cao et al., 2021). In addition, earlier species delineation based on macroscopic and microscopic morphology overestimated the range of distribution of some “species” because this concept did not account for other types of differentiation. For example, spore morphology and macroscopic features of sporocarps are effectively identical between some *Pleurotus* species, yet host preference, environmental differences, sexual incompatibility, and genetic markers set these species apart (Li et al., 2017; Segedin et al., 1995; Vilgalys et al., 1993). The advent of the phylogenetic species concept and molecular tools has led to the discovery of numerous cryptic species, splitting existing species that were previously delineated by morphology and ecology into multiple species delineated only by genetic markers (Boekhout et al., 2021; Hughes & Petersen, 2015). In recent years, the combination of multiple genetic markers with complex statistical phylogenetic analyses has enabled delineation of distinct species, even within genera that display remarkable variation in the most commonly used marker for molecular species delimitation in fungi, the internal transcribed spacer (ITS) region of the ribosomal DNA (Chethana et al., 2021; Vizzini et al., 2024). Genes are known to evolve faster than morphological features (Taylor et al., 2006), which might explain the discovery of many cryptic species. Multi-locus phylogenetics coupled with morphology and, in some cases, whole genome phylogenetics, has become the gold standard for describing a species in fungal taxonomy (Aime et al., 2021; Cao et al., 2021; Cazabonne et al., 2024; Xu, 2020). Nevertheless, there is still no definitive consensus on defining a fungal species. In *Pleurotus*, interfertility is possible between sibling species (Menolli et al., 2014; Petersen & Ridley, 1996; Rosnina et al., 2016), further blurring species boundaries. More recently, molecular sequencing used in combination with morphological descriptions has helped clarify species delimitation within the genus, but most published studies to date have not included *Pleurotus* species present in New Zealand, or only a subset of them (Li et al., 2020; Pánek et al., 2019; Vizzini et al., 2024; Zervakis et al., 2019).

The three main species indigenous to New Zealand are *Pleurotus australis* (Cooke & Massee) Sacc.*, P. parsonsiae* G. Stev. and *P. purpureo-olivaceus* (G. Stev.) Segedin et al., all three of which are also present in Australia (according to iNaturalist, https://www.inaturalist.nz). In addition, *P. abalonus* Y.H. Han et al. is indigenous to Raoul Island, an island far north of mainland New Zealand in the Pacific Ocean (PDD 113796; collection from “PDD”, the New Zealand Fungarium, Te Kohinga Hekaheka o Aotearoa). There is one isolated record of *P. velatus* Segedin et al. known only from the type specimen in Waitakere Ranges Regional Park, Auckland (Segedin et al., 1995), which may have been a misidentification.

Previous studies have suggested that both *P. australis* (Petersen et al., 1997) and *P. purpureo-olivaceus* (Petersen, 1992; Segedin et al., 1995) have anamorphic stages, which are rare in *Pleurotus* and are confined to the subgenus *Coremiopleurotus* (Zervakis et al., 2004). However, the placement of *P. purpureo-olivaceus* within *Coremiopleurotus* by Segedin et al. warrants further investigation, because evidence of an anamorphic stage of this species from two historic records (Petersen, 1992; Segedin et al., 1995) is not supported by contemporary observations.

It is unclear from existing literature whether *P. pulmonarius* (also known as grey oyster mushroom or phoenix oyster mushroom), a species that is both naturally occurring and cultivated in New Zealand, is introduced or indigenous (Segedin et al., 1995). The first record of importation for cultivation was from 1994 (Buchanan & Barnes, 2002; Wassilieff, 2008), but wild collections predate this as far back as 1980 (PDD 91221). Petersen & Ridley (1996) described a “New Zealand *P. pulmonarius*” strain (NZFRI3528) that was mating-compatible with international isolates of *P. pulmonarius*, *P. ostreatus*, *P. eryngii* and *P. abieticola* R.H. Petersen & K.W. Hughes, all of which fall under the larger *P. ostreatus* complex (Li et al., 2020; Pánek et al., 2019). Interestingly, other international *P. pulmonarius* isolates are incompatible with *P. ostreatus* (Petersen & Hughes, 1993). Thus, “New Zealand *P. pulmonarius*” behaves differently than international *P. pulmonarius*. No further research has been conducted on *P. pulmonarius* in New Zealand, but several local mushroom cultivators claim that their wild strains are different from international ones (Acres, 2024).

Aside from these four species found throughout New Zealand (Figure 1), the tropical *P. djamor* (Rumph. ex Fr.) Boedijn (commonly known as pink oyster mushroom) was imported and cultivated by small-scale growers. Many of these growers offer inoculum and home growing kits for indoor and outdoor cultivation. *P. djamor* is closely related to *P. parsonsiae* and *P. opuntiae* (Durieu & Lév.) Sacc., with the species names used interchangeably in some previous publications (Segedin et al., 1995; Segedin & Pennycook, 2001; Zervakis et al., 2019). The import of new species into New Zealand is generally prohibited by the *Hazardous Substances and New Organisms Act 1996.* In 2015, the Environmental Protection Authority of New Zealand approved *P. djamor* for importation on the grounds that the species was recorded as present in New Zealand prior to 1996 and therefore not considered to be a ‘New Organism’ under the terms of the Act. That decision was based on the historical evidence for the presence of *P. opuntiae* (*HSNO Application PNZ1000218 | EPA*, 2015). However, the name *P. opuntiae* in New Zealand was misapplied and based on specimens that were later identified through DNA sequencing to be a distinct indigenous species *P. parsonsiae*, not *P. djamor* (Zervakis et al., 2019).

**Figure 1:**
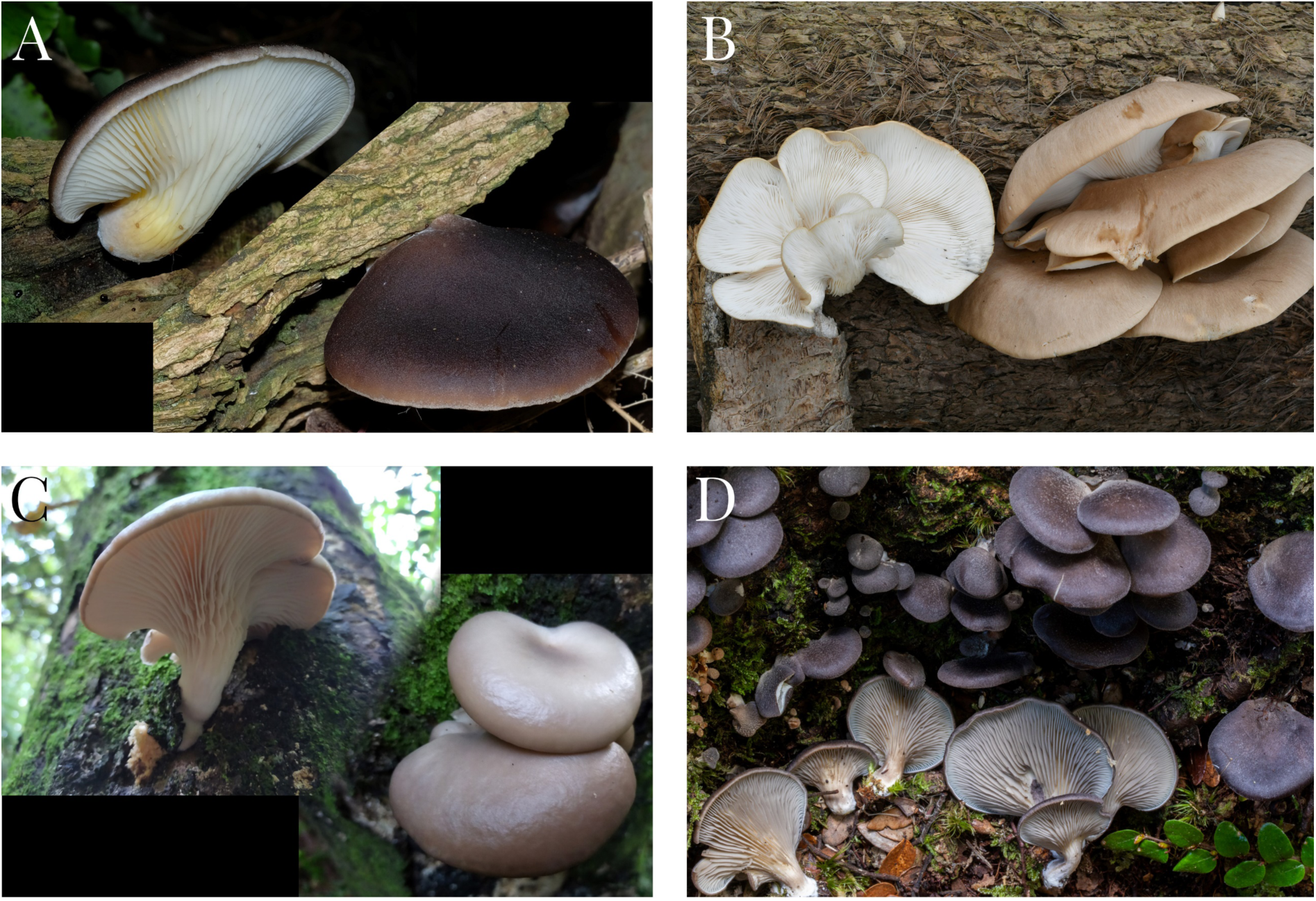
Four major species of Pleurotus in New Zealand. Mature sporocarps of Pleurotus australis (A), P. parsonsiae (B), P. pulmonarius (C) and P. purpureo-olivaceus (D) from different locations. Photo credit, all CC-BY-NC: Aman Hunt (A; iNaturalist #102978764), Christian Schwarz (B; iNaturalist #25051299), Barton Acres (C; iNaturalist #259097346), and Dean Lyons (D; iNaturalist #209189504).

This study elucidates the molecular phylogeny of indigenous and exotic *Pleurotus* species in New Zealand, using existing and new collections from the wild, and from mushroom growers, for DNA sequencing and a multi-gene phylogenetic analysis of five rDNA loci. We test the hypothesis that indigenous *P. pulmonarius* strains are genetically and morphologically distinct from imported international strains and propose a name for a new indigenous variety of *P. pulmonarius.* We further investigate the evidence against an anamorphic stage of *P. purpureo-olivaceus*. Our results highlight the potential for using indigenous oyster mushrooms for sustainable and viable cultivation in New Zealand.

## Methods

What follows are the condensed methods, with the extended methods available in the supplement.

### Biological material

We collected 84 *Pleurotus* specimens from New Zealand for this study (Table A1). 43 specimens were sampled from the wild in 2021 and 2022, 19 strains were sourced from International Collection of Microorganisms from Plants (ICMP) and 22 strains from commercial suppliers. Pure cultures and dried vouchers were prepared for each specimen.

To investigate evidence of an anamorphic stage of *P. purpureo-olivaceus*, we macro- and microscopically examined the type collection and vouchers collected by Segedin et al. (1995). We further examined the culture ICMP 9629 described by Petersen (1992), as well as all specimens observed in the wild and in culture.

### Taxonomic description

Dried material of *P. pulmonarius* specimens was soaked in 5% KOH before sectioning and microscopic examination. Measurements were taken from fruitbodies collected in the wild, and from cultivation of both wild-type and imported strains.

### DNA extraction, PCR amplification, sequencing and data assembly

We generated biomass for DNA extraction from liquid cultures and followed a manual cetyltrimethylammonium bromide (CTAB) based DNA extraction protocol (adapted from Jones & Schwessinger, 2021) to generate high molecular weight DNA for Sanger sequencing and short-read whole genome sequencing. DNA extraction from nine samples failed to produce sufficient quality DNA for subsequent steps, dropping the total number of samples for sequencing to 75.

To investigate the phylogenetic relationships among *Pleurotus* samples, we analysed five molecular markers: the internal transcribed spacer region (“ITS” here, referring to ITS1+5.8S+ITS2) and the large subunit (LSU) of the nuclear ribosomal RNA gene, the first (RPB1) and second subunits of RNA polymerase II (RPB2), and the translation elongation factor alpha (Tef). The ITS region is often used for species identification of fungi but can be inconclusive for the phylogenetic distinction of closely related species in *Agaricales* (Davison et al., 2017; Klopfenstein et al., 2017). ITS sequences show significant degrees of intraspecific variability, which may lead to errors in species delimitation (Estensmo et al., 2021; Hilário et al., 2022; Nilsson et al., 2008; Tremble et al., 2020; Wilson et al., 2023). Two studies of *Pleurotus* found that Tef and RPB2 provided sufficient resolution to distinguish different species within the *Pleurotus eryngii* species complex (Estrada et al., 2010; Zhao et al., 2016).

From the DNA extracts, only ITS, RPB2 and Tef were amplified directly, whereas LSU and RPB1 were *in silico* extracted from whole genome assemblies as a complementary approach to add additional targets. PCR amplification details are outlined in the Extended Methods. PCR products were sent to Ecogene (Auckland, New Zealand) for bidirectional Sanger sequencing. We aligned and trimmed reverse and forward sequences using Geneious 10.2.6 (Biomatters, Inc., New Zealand) and Sequencher 5.4.6 (Gene Codes Corporation, USA).

### Library preparation, sequencing and *de novo* genome assembly

In addition to Sanger sequencing, we prepared 54 samples for short-read whole genome sequencing using the MGI platform (MGI Tech, China), for subsequent *in silico* amplicon extraction. 30 samples were excluded from genome sequencing. 150-bp paired-end sequencing of the DNA library was performed using DNBSEQ-G400 (MGI Tech, Beijing, China) by a commercial sequencing service (Lincoln University, New Zealand).

Demultiplexed reads were quality trimmed to Q30 using BBMap v39.01 (Bushnell, 2023) and *de novo* assembled using SPAdes v4.0.0 (Bankevich et al., 2012) with default parameters.

### *In silico* target gene region extraction from genomes

Some key samples did not amplify well for ITS, RPB2 and/or Tef using Sanger sequencing with standard primers and PCR conditions; therefore, we used two different methods to *in silico* extract ITS, RPB2 and Tef from the *de novo* assembled genomes, with the addition of LSU and RPB1.

First, we utilised a BLASTn subject query (Camacho et al., 2009) to extract matches of representative regions of ITS, LSU, RPB1, RPB2, and Tef within each genome assembly using closely related reference sequences from GenBank for each species, where available. Second, we used ThermonucleotideBLAST v2.61 (“tntblast”, Gans & Wolinsky, 2008) to extract predicted amplicons from each genome assembly using forward and reverse primers for each target region. The BLASTn approach had a higher success rate but resulted in shorter amplicons than tntblast (Table B3).

To further strengthen the multi-gene phylogenetic analysis, we extracted the same loci from other *Pleurotus* reference genomes on NCBI, as well as several other closely related *Agaricales* taxa genomes (Table A2), in order to have full representation of the same loci across outgroups and ingroups.

### Sequence alignment and phylogenetic analyses

We chose the *in silico* extracted amplicons for subsequent steps, supplementing them with amplicons from taxa which only Sanger sequencing data was available for. The genome-extracted amplicons were given preference due to the higher quality of reads and better representation of the target regions (Table B3), especially RPB2, for which only partial gene amplification was successful with Sanger sequencing using a single primer pair.

Additional sequences from ICMP and PDD were included, along with ITS, LSU, RPB1, RPB2 and Tef sequences of other taxa from GenBank. No international *Pleurotus* sequences in GenBank that were related to the New Zealand species had sufficient representation for all five gene regions.

Sequences were aligned separately for each region using MAFFT v7.450 and the E-INS-i algorithm (Katoh & Standley, 2013). Alignments were manually realigned in Geneious, to correct misaligned gaps across related taxa. A multi-gene alignment was generated by concatenating the five amplicon alignments.

Phylogenetic analysis was based on maximum likelihood and Bayesian inference for each individual gene region and the concatenated alignment. Maximum likelihood (ML) analyses were conducted with IQ-TREE v2.3.6 (Nguyen et al., 2015), using the -MFP command to automatically choose the appropriate substitution model for each partition (using the -p command). Bootstrap values were computed using Ultrafast analysis of 1000 pseudoreplicates, to evaluate branch confidence and find the best-scoring ML tree. Bayesian phylogenies were constructed using MrBayes v3.2.7a (Ronquist et al., 2012), using the optimal substitution models as identified by IQ-TREE ModelFinder for each partition. Posterior probabilities were estimated using common settings (details see Extended Methods), with four million generations.

Clade support was considered significant when thresholds of > 90% for bootstrap support and > 95% for posterior probability were used (Alfaro et al., 2003). Trees were visualised using Geneious and further annotated using InkScape v1.3.2 (https://inkscape.org). *Sarcomyxa edulis*, *Pterula gracilis*, *Typhula micans* and *Hygrocybe conica* were used as outgroups in all trees.

Initial phylogenetic analyses included all newly sequenced taxa (Table A1), but were filtered for the final analysis by removing highly similar sequences (> 99.5% sequence identity) and to balance out the number of sequences per taxa. In total, 57 taxa were used in the final analysis (Table A2).

### Distribution mapping

We constructed a distribution map of *Pleurotus* in New Zealand using QGIS 3.34 (QGIS Association) with metadata from specimens held at PDD and cultures at ICMP. We further included genus-level observations from iNaturalist (https://www.inaturalist.nz, accessed 16/07/2024), as well as “Research Grade” observations, which refers to species-level observations with a majority consensus identification by the community.

### Molecular species delimitation

To test the hypothesis of a distinct indigenous *P. pulmonarius* clade, we investigated the sequence similarity for each locus. First, we generated a consensus sequence of all strains in each clade identified in the phylogenetic analysis, removing any indels. The two consensus sequences were then aligned with MAFFT v7.450 using the E-INS-i algorithm. MAFFT calculated a pairwise sequence similarity score, which we then used to assess speciation. For many genera of fungi, a threshold of 97-98.5% sequence similarity in ITS is used to differentiate species (Lücking et al., 2020).

Furthermore, we used the species delimitation plugin v1.4.5 (Masters et al., 2011) in Geneious to calculate the following statistical metrics: (i) *P_RD_* (Randomly Distinct), which represents the probability of clades being distinct as a result of random coalescent processes; (ii) Rosenberg’s *P_AB_*, which expresses the probability of reciprocal monophyly of the clade of interest and its nearest defined group under random branching; and (iii) *P_ID_* (Liberal), which is the mean probability (with 95% confidence intervals) of correctly identifying a clade as a species based on the difference between the intra- and interspecific genetic distances.

## Results

### Distribution of *Pleurotus* in New Zealand

As of July 2024, there were 180 fungarium collections (PDD; excluding new vouchers from this study), 43 culture cultures (ICMP; excluding new cultures from this study), and 483 “Research Grade” observations on iNaturalist of *Pleurotus* within New Zealand. Additional iNaturalist observations could only be identified to the genus level based on the details and photos provided.

Each *Pleurotus* species has a distinct distribution across New Zealand (Figure 2). *P. pulmonarius* is widespread across the North and South Island, often observed near urban centres. Most genus-level iNaturalist observations are likely attributable to *P. parsonsiae* or *P. pulmonarius* because these two species are often difficult to identify from photos. *P. parsonsiae* and *P. australis* are more common on the North Island than the South Island, while *P. purpureo-olivaceus* is more common in the South Island. The distribution of *P. purpureo-olivaceus* closely follows that of southern beech trees (*Nothofagus* spp.), its indigenous hosts. However, *P. purpureo-olivaceus* has also been observed on exotic trees such as birch (*Betula* spp.), gorse (*Ulex* spp.) and willow (*Salix* spp.). *P. australis* is most abundant in the upper half of the North Island, and sporocarps of *P. australis* have been collected and observed from living indigenous and exotic trees (PDD 120236, iNaturalist #101389646, iNaturalist #197760600, iNaturalist #110440505), as well as from dead wood.

**Figure 2:**
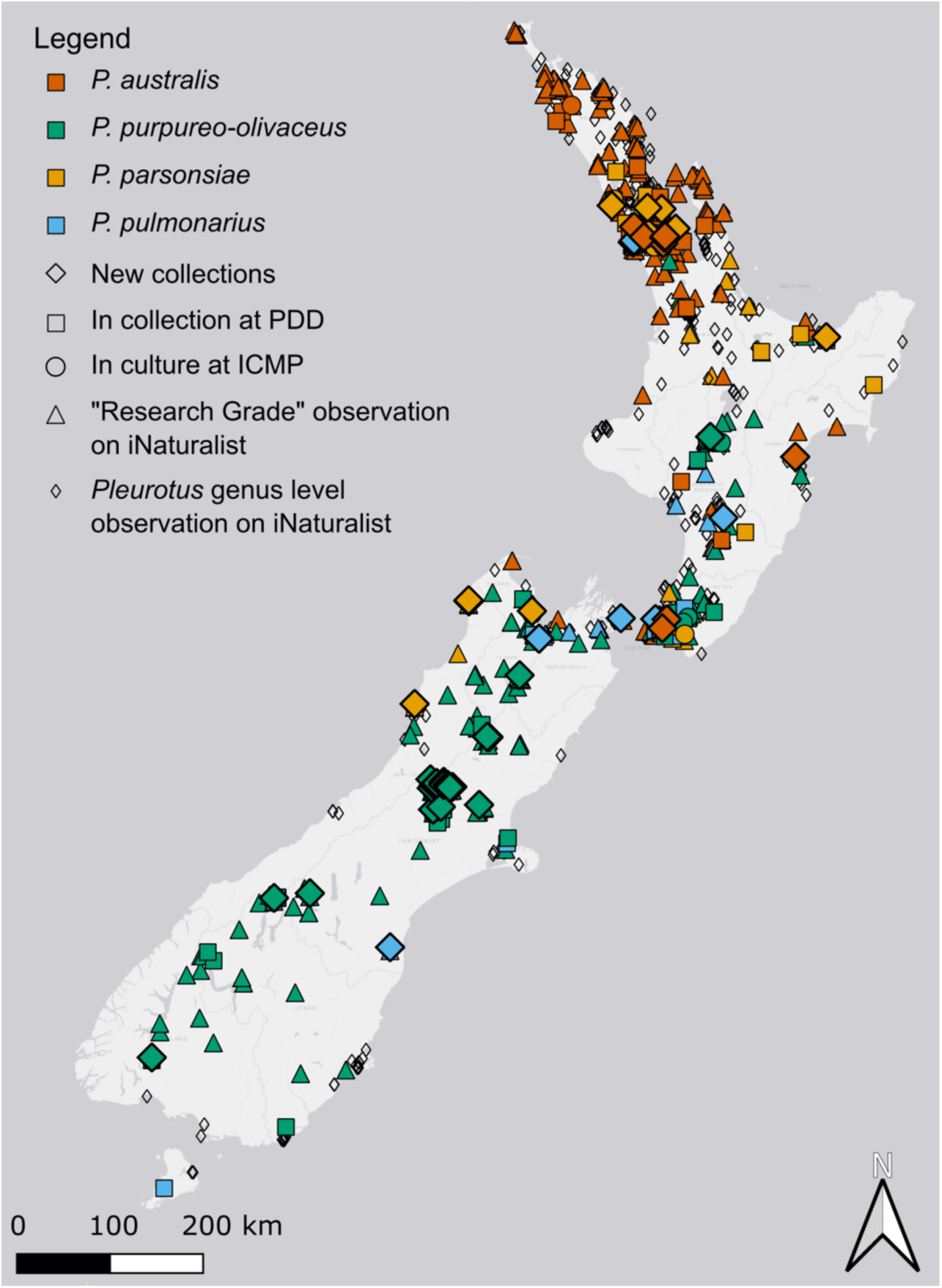
Distribution map of Pleurotus records in Aotearoa. Data sources: New Zealand Fungarium, Te Kohinga Hekaheka o Aotearoa (PDD), International Collection of Microorganisms from Plants (ICMP), and iNaturalist (https://www.inaturalist.nz/). Sources are marked by filled diamonds (collections from the present study), squares (PDD), circles (ICMP), triangles (“Research Grade” iNaturalist observations), and open diamonds (genus level iNaturalist observations). Species are marked by colour: P. australis (brown), P. parsonsiae (tan), P. pulmonarius (blue) and P. purpureo-olivaceus (green). All records are dated: 16/07/2024

### Phylogenetic analysis

The final dataset comprised 57 taxa, including both newly sequenced specimens and publicly available sequences from GenBank. The multi-gene phylogenetic analysis of New Zealand *Pleurotus* using ITS, LSU, RPB1, RPB2 and Tef (Figure 3) showed identical topology in both the maximum likelihood analysis and Bayesian inference, with strong support for the early diverging taxa *P. purpureo-olivaceus* and *P. tuber-regium* (>95% bootstrap support, 1.0 posterior probabilities). The positions of *P. australis* and the other two international taxa in the *Coremiopleurotus* clade were strongly supported (>95% bootstrap support, 1.0 posterior probabilities), with a significant support for the position of the *Coremiopleurotus* clade in relation to the clade containing *P. djamor* in the Bayesian inference (1.0 posterior probability) but not in the maximum likelihood analysis (81% bootstrap support). Both of these clades had strong support in their separation from the *P. ostreatus, P. eryngii, P. pulmonarius* clade (93% bootstrap support, 1.0 posterior probabilities), with strong interior branch support within each of these clades (>99% bootstrap support, 1.0 posterior probabilities). Each named species was clearly separated from the other species, supporting the monophyletic origin of all taxa in this study. The single-gene trees were unable to resolve the relationships between the studied taxa (weak to moderate statistical support), and showed variable topology compared to the multi-gene tree and to trees from other gene regions (Figures B1-B5).

**Figure 3:**
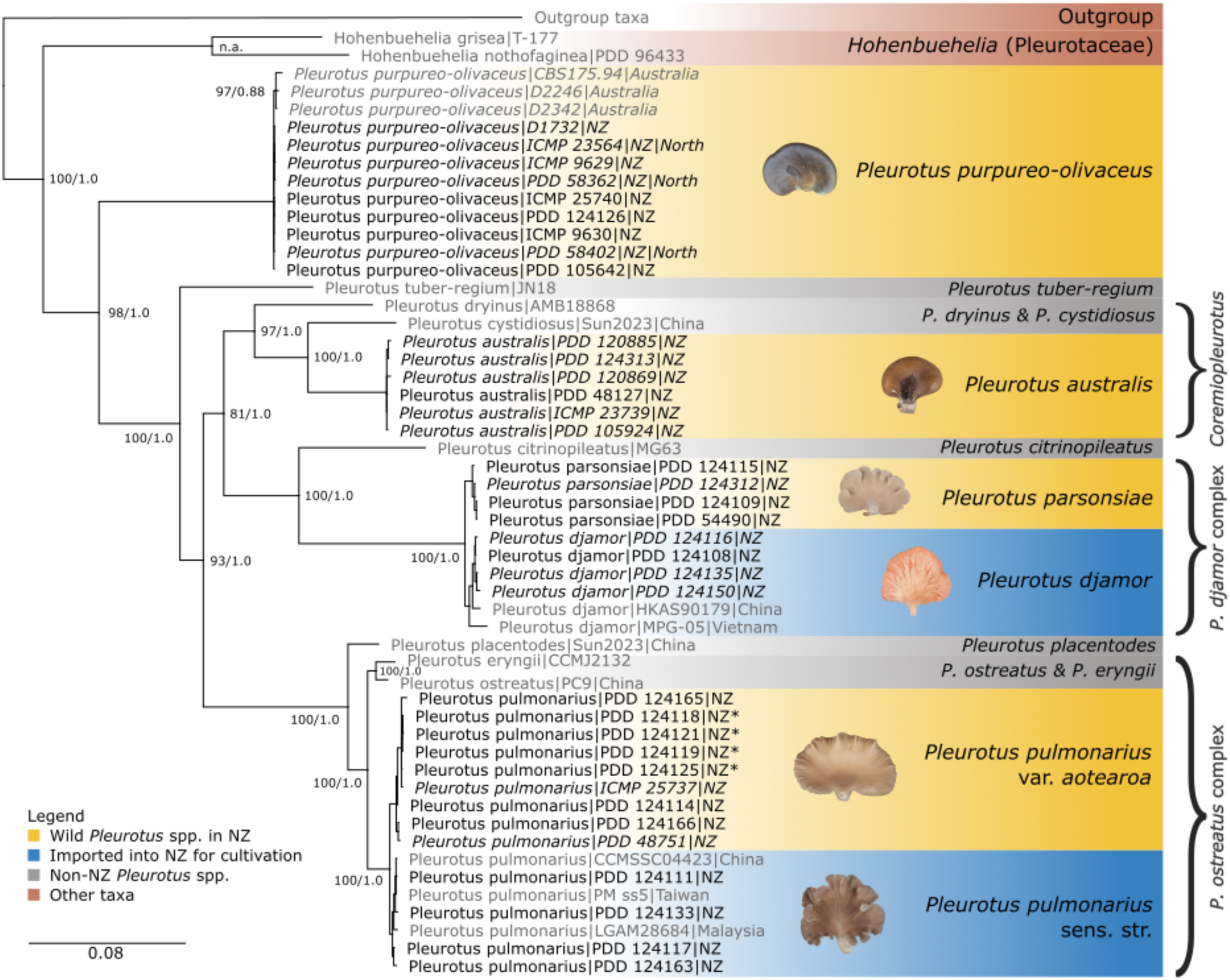
Multi-gene phylogenetic tree of New Zealand (NZ) Pleurotus, using maximum likelihood and Bayesian inference of concatenated ITS-LSU-RPB1-RPB2-Tef alignment. Numbers next to nodes denote bootstrap support/posterior probabilities. Taxa not found in NZ are marked in grey. Sequences from wild NZ collections are highlighted in yellow and imported taxa are highlighted in blue. Cultivated P. pulmonarius taxa isolated from the wild are marked with a star. Hohenbuehelia sequences were included to inform the position of P. purpureo-olivaceus within Pleurotus. The outgroup taxa (collapsed here for graphical representation) were Hygrocybe conica, Pterula gracilis, Sarcomyxa edulis and Typhula micans. Taxa in italics are missing sequencing data in one or more gene regions. Clades belonging to the subgenus Coremiopleurotus are marked with a vertical bar. All NZ P. purpureo-olivaceus strains are from the South Island, except those marked “North” for North Island collections.

### Support for an indigenous *P. pulmonarius* variety

We identified a distinct clade corresponding to *P. pulmonarius* var. *aotearoa* specimens, containing samples originating from the wild and cultures from growers claiming a wild origin. The concatenated dataset strongly supported a clear split between *P. pulmonarius* var. *aotearoa* and *P. pulmonarius sens. str.* (100% bootstrap support, 1.0 posterior probabilities), which were also significantly separated in the individual RPB1, Tef, and RPB2 phylogenies, the latter however only in the Bayesian inference. *P. pulmonarius* appeared to be polyphyletic in the ITS analysis, with 99.9% sequence similarity between clades. Sequence similarities between the two clades were also high for LSU (99.9%), but lower for the other loci (RPB1 97.4%, RPB2 98.0%, Tef 94.9%).

In the multi-gene phylogeny, both the maximum likelihood tree analysis and Bayesian inference indicated speciation (Rosenberg’s *P_AB_* < 0.01; Table 1). The distance between the two *P. pulmonarius* clades was in the same order of magnitude as the distance between *P. parsonsiae* and *P. djamor* in the Bayesian inference (0.02 vs. 0.03 average pairwise tree distance, respectively). The mean probability of correctly identifying *P. pulmonarius* var*. aotearoa* as a species was 0.94 (*P_ID_* (Liberal) confidence intervals between 0.86-0.99). A non-significant result in the *P_RD_* values inferred in the Bayesian phylogeny of *P. pulmonarius* var. *aotearoa* (*P_RD_* > 0.05; Table 1) was contrasted with a significant result in *P. pulmonarius* sens. str. (*P_RD_* = 0.05). The same metric was not significant in *P. djamor* or *P. parsonsiae* in the Bayesian approach either (*P_RD_* > 0.05), but was significant in all the maximum likelihood analyses.

**Table 1:**
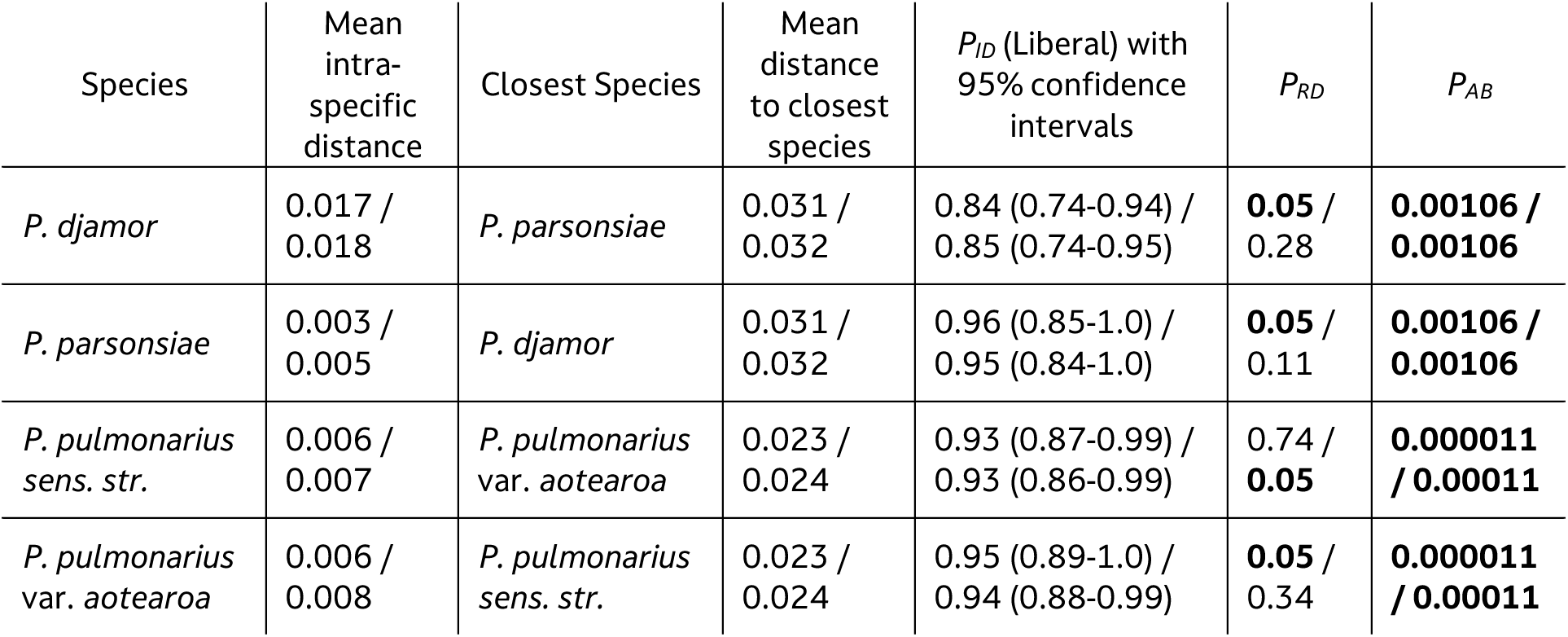
Summary of the species delimitation statistics for two clades (P. djamor, P. parsonsiae, and P. pulmonarius var. aotearoa and sens. str.) as calculated from maximum likelihood (left values) and Bayesian inference trees (right values). P values of P_RD_ (Randomly Distinct) and Rosenberg’s P_AB_ in bold are statistically significant at P ≤ 0.05. Mean intraspecific distance and distance to closest species are average pairwise tree distances.

### Taxonomic description

*Pleurotus pulmonarius* var*. aotearoa* D. Hera & J.A. Cooper var. nov. Holotype PDD 124119, DH1337-O (iNat259097346) MB: ####

### Macroscopic features

**Pileus:** diam. 20 - (80) - 150 mm (σ = 2.9), spathulate, rounded, flabelliform to reniform, depressed above the laterally attached stipe. **Pileus edge** entire to weakly incised, sometimes lobed, planar to weakly crenulate, and weakly striate at the perimeter when moist. **Colour** greyish-orange (6B3)

**Lamellae:** concolorous with the pileus, crowded, thin, up to 5 mm deep, mixed in with regularly occurring lamellulae of varying length. Lamellae long-arched, running into the stipe base as narrow ribs and usually reaching near to the stipe base, not anastomosing at the base but with occasional bifurcation. Surface white, ivory colored (4B3), to cream (4A3), faint yellowing when bruised. **Lamellar edge** concolorous, smooth, sometimes notched towards the stipe base.

**Stipe:** eccentric to lateral, cylindrical, base strigose with coarse-concolorous fibres. Stipe 3 - (15) - 36 mm diam. (σ = 0.6) × 6 - (14) - 24 mm long (σ = 0.5). Stipe length/pileus diam. ratio 0.2 (σ = 0.12). **Stipe flesh** compact, white, weakly yellowing

**Taste:** Mild, pleasant.

**Odour:** neutral.

**Microscopic features**

**Spores:** (Figure 4C) smooth, hyaline, inamyloid, subcylindrical, with a lateral apiculus. Holotype: length 10.1 µm (σ = 0.64), width 4.4 µm (σ = 0.28), Q = 2.31 (σ = 0.12), n = 20. Combined holotype + paratypes (PDD 124119, PDD 124114, PDD 124165, PDD 124166): length 10.2 µm (σ = 0.67), width 4.7 µm (σ = 0.26), Q = 2.2 (σ = 0.15), n = 20+20+20+20. Spores frequently germinating in-situ.

**Figure 4:**
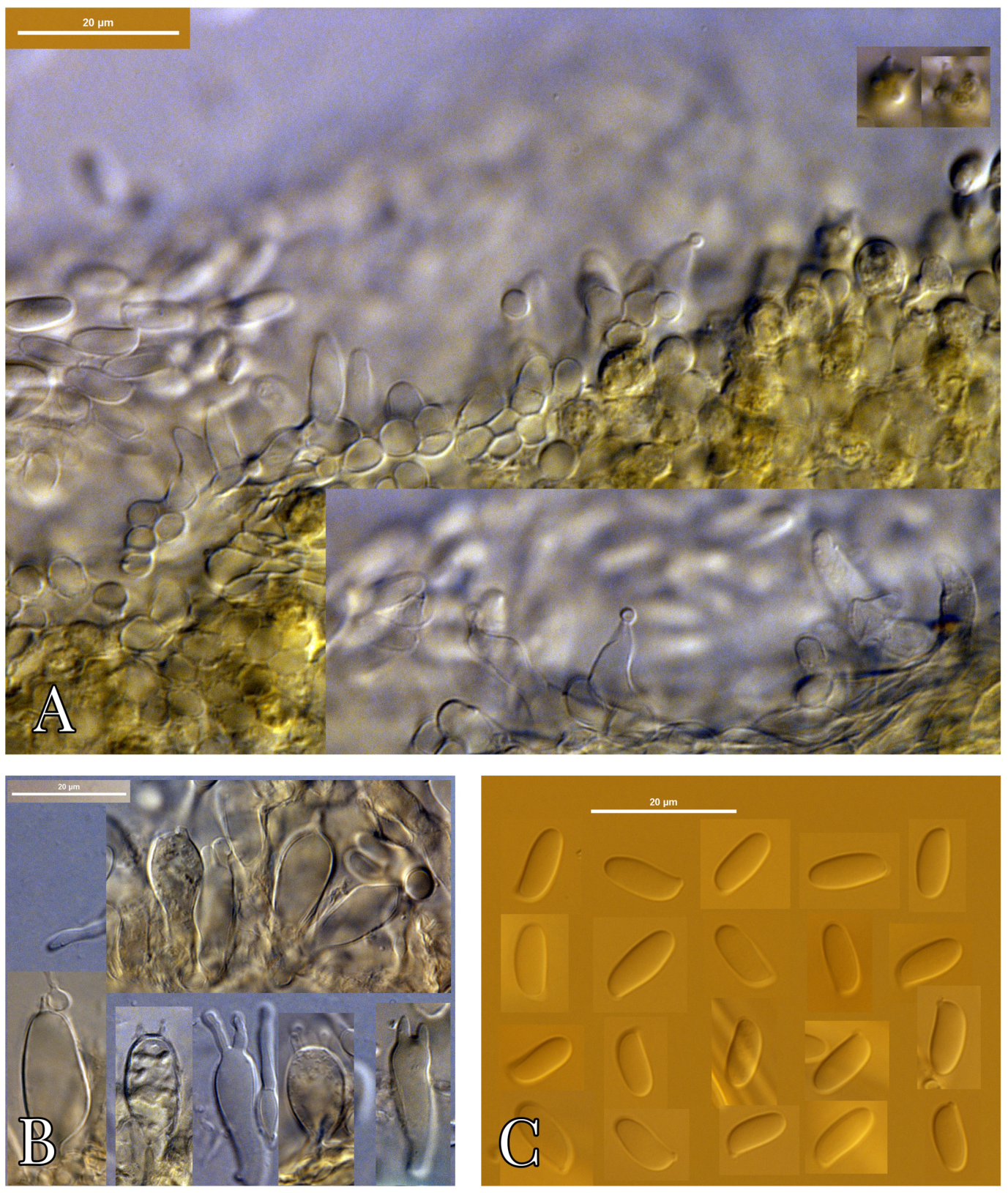
P. pulmonarius var. aotearoa (Holotype, PDD 124119) micro morphological features in 5% KOH. (A) lamellar edge showing cheilocystidia; (B) basidia; (C) spores.

**Basidia:** (Figure 4B) 15-30 × 6 – 11 µm, subcylindrical to clavate, with basal clamp, sometimes with yellow oil drops, mostly 4-sterigmate (infrequently 2 or 3 sterigmate). Sterigma to 5 µm. long. **Basidioles:** subcylindrical.

**Pleurocystidia:** not observed. Hyphal pegs (Hilber, 1982) not observed.

Lamellar edge fertile, with infrequent cylindrical to geniculate **cheilocystidia** (Figure 4A) to 15 × 5 µm, frequently with a mucronate apex.

**Caulocystidia:** not observed

**Pileipellis:** an interwoven cutis of clamped, refractive-walled hyphae (but not sclerified), 5 – 8 µm diam., without conspicuous terminal elements. Cutis to 150 µm deep. No oleiferous hyphae observed.

**Subhymenium & trama** hyphae to 7 µm diam. Less densely packed than the cutis. Occasional weakly sclerified hyphae observed.

Lamellar trama: irregular, loosely woven, of thin-walled hyphae, 3 – 12 µm diam. Oleiferous hyphae absent.

**Stipe cortex:** monomitic, consisting of parallel, densely packed, clamped, refractive-walled hyphae, to 4 µm diam. Occasional sclerified hyphae observed.

**Stipe hairs:** tightly packed agglutinate bundles of hyphae 20 – 40 µm in diam.

### Ecology

**Habitat:** solitary to imbricate, bases never fused. Frequent on *Cordyline* but also *Sophora*, *Hoheria*, and introduced *Cassia* and *Acacia*.

**Distribution:** Throughout New Zealand but more common in the north on cabbage tree and angiosperms.

**Notes:**

*P. pulmonarius* was originally described from Sweden from fruitbodies associated with Birch trees. Hilber (1982) revised the European species of *Pleurotus* and studied the morphology of *P. pulmonarius* fruitbodies from various host trees. The species does not have a modern interpretive epitype. The origin of commercial strains in cultivation in New Zealand is mostly unknown but phylogenetic data indicate an Asian origin.

Collections of *P. pulmonarius* from indigenous habitats in New Zealand possess some consistent morphological differences to fruitbodies derived from imported commercial strains, and from Hilber’s concept of the European wild type. In New Zealand fruitbodies from the wild, or from ex-wild type strains, the stipe is relatively short, the lamellae relatively deep, and they run almost the entire stipe length without a significant decrease in depth, and there is no anastomosing between the lamellae at the stipe base. Microscopically the spores are relatively large and frequently > 10 µm in length. In some specimens there is abundant germination of spores *in situ*. Fruitbodies from cultivated imported strains and Hilber’s description of European wild-type have relatively long stipes, the lamellae terminate well before the stipe base, become shallow, and with clear anastomosing between lamellae. Microscopically the spores of *P. pulmonarius sens. str.* are generally shorter and often < 9 µm long. The combined spore data for examined non-New Zealand specimens (PDD 124110, 124112 and 124160) are: length 8.7 µm, (σ = 0.44), width 4.1 µm (σ = 0.23), Q = 2.15 (σ = 0.14), n = 17+20+20.

In New Zealand *P. pulmonarius* var. *aotearoa* may be distinguished from the similarly coloured dimitic *P. parsonsiae* because the latter has a much tougher consistency due to the presence of abundant sclerified hyphae, and the fruitbodies are often fused at the base. *P. parsonsiae* lamellae are also more narrowly spaced than *P. pulmonarius* var. *aotearoa* lamellae.

Although a rare occurrence, *P. pulmonarius* var. *aotearoa* has been observed to fruit on living trees (iNaturalist #26320495 on *Cordyline*). This phenomenon is also seen in *P. australis* (PDD 120236, iNaturalist #101389646, iNaturalist #197760600, iNaturalist #110440505). Several species within the *P. eryngii* complex also grow on living plants (Jia et al., 2024), but it remains unresolved whether they act as pathogens or grow saprotrophically on necrotic plant tissue (Carlavilla & Manjón, 2023). Further research is required to determine the lifestyle of *P. pulmonarius* var. *aotearoa* and *P. australis*.

**New Zealand specimens examined:** NEW ZEALAND: SOUTH ISLAND: DUNEDIN—Outram Glen Scenic Reserve, on *Hoheria* sp., *B. Acres*, 11 Jun. 2021, PDD 124119 (**holotype** of *P. pulmonarius* var. *aotearoa*). NELSON—Wairoa Gorge, on *Cordyline australis*, *J. Harding*, 14 Feb. 2022, PDD 124165 and PDD 124166. SOUTH ISLAND: MARLBOROUGH—Arapaoa Island, on *Cordyline australis*, *T. Hitchcock*, 9 May 2019, PDD 124114.

**Imported specimens examined:** The following specimens were of unknown origin outside of New Zealand, but sourced and cultivated from commercial grow kits. Supplier names are in italics. *Mushroom Gourmet,* Whangarei, isolated 24 Feb. 2022, PDD 124110; *Urban Fresh Farms*, Masterton, isolated 24 Feb. 2022, PDD 124112; *Oak & Spore*, Christchurch, isolated 8 Jun. 2022, PDD 124160.

#### P. purpureo-olivaceus examination

Examination of *P. purpureo-olivaceus* specimens in the wild (52 of our own observations and photos of a total of 222 observations on iNaturalist) and 49 cultures showed no evidence of developing an anamorphic coremial state. The revived culture ICMP 9629, described by Petersen (1992) as producing arthrospores, showed only typical mycelial growth. Unfortunately, the putative “*P. purpureo-olivaceus* anamorph” vouchers (PDD 61128 and 61129) described by Segedin et al. (1995) could not be located in PDD. Furthermore, there was no evidence of asexual spores in other collections from Segedin et al. collected around the same time (PDD 44961, 58355, 58360, 58362, 58402), or in the type specimen of *P. purpureo-olivaceus*.

## Discussion

### Multi-gene phylogeny shows clear separation of *Pleurotus* in New Zealand

New Zealand is home to four distinct species of *Pleurotus*, from the earliest diverging lineage in the genus, *P. purpureo-olivaceus*, to *P. parsonsiae* as the most derived species. The multi-gene tree topology presented here matched that of two recent phylogenetic analyses in the Agaricales using six-gene (Vizzini et al., 2024) and 40-gene (Li et al., 2020) datasets, as well as one analysis based on ITS only (Menolli et al., 2014). However, a single-gene phylogenetic analysis by Zervakis et al. (2019), which included several New Zealand and overseas *Pleurotus* species, had a different topology, with *P. australis* in a sister relationship with a larger clade containing *P. parsonsiae* and *P. ostreatus*. The contrasting result from Zervakis et al. might be related to the reliance on a single gene, as it is the same as our ML and Bayesian ITS trees (Figure B1). We found low utility using single-gene phylogenies (Figures B1-B5) to resolve *Pleurotus*. Our results indicate that a five-gene dataset was required to confidently describe coarse topology within the genus. We found that all *Pleurotus* species naturally present in New Zealand were monophyletic.

### Low genetic diversity in Pleurotus purpureo-olivaceus

*P. purpureo-olivaceus* is of special interest because it is a separate lineage to all other known *Pleurotus* species and showed minimal genetic differentiation across populations in New Zealand. A recent phylogenomic study of Basidiomycota estimated a divergence time of 73 million years for *P. tuber-regium* from the most recent common ancestor (He et al., 2024), which would place *P. purpureo-olivaceus* (which was not included in that study) even earlier. The New Zealand *P. purpureo-olivaceus* sequences were highly similar (> 99.5% mean pairwise sequence identity, from MAFFT alignment distance matrices for each locus of all 32 strains of *P. purpureo-olivaceus*), so we excluded the majority of sequences from phylogenetic analysis based on redundancy.

To date, GenBank accessions of international *P. purpureo-olivaceus* specimens are limited to a few ITS, LSU and Tef sequences. However, the reliability of some GenBank sequence metadata is questionable, including country origins and species assignments. For example, ITS and Tef sequences identical to New Zealand *P. purpureo-olivaceus* strains have been mislabelled as *P. eryngii* and *P. fossulatus*, and there are spurious collection records from India, South Korea and Czechia. We also found instances of a single voucher number showing inconsistent species assignments and collection countries across different GenBank sequence submissions, e.g. with specimen D2246 (Table A2).

This is further confounded by the same voucher number having different metadata depending on the sequence region accessioned. For example, the country of the ITS sequence of “*P. rattenburyi*” (a synonym of *P. purpureo-olivaceus*) isolate P103 is recorded as South Korea in GenBank (MG282502), but its original collection number KACC #46280 (Lin et al., 2022) matches CBS 175.94 on the KACC website (Korean Agricultural Culture Collection). CBS 175.94 was indeed collected from Tasmania and is also in GenBank as a separate accession with no country metadata (EU424315). This example highlights common data quality issues in GenBank (Hofstetter et al., 2019), where in fact the same voucher has two ITS sequences accessioned, neither with the correct country metadata. Thus, we conclude that GenBank records of *P. purpureo-olivaceus* outside of Oceania are spurious.

Some of these GenBank sequences were able to be linked to Australian specimens, which are only differentiated from New Zealand specimens in ITS (98.4–99.2% sequence similarity between Australian and New Zealand taxa, compared to 99.7–100% sequence similarity within New Zealand taxa), but not in LSU or Tef (see also supplement figures of single gene trees). While these findings indicate that Australian *P. purpureo-olivaceus* populations share a tendency for low genetic diversity with New Zealand populations, this would need to be ascertained in future studies, requiring the sequencing of more specimens from Australia using all five loci or whole genomes.

### *P. pulmonarius* var. *aotearoa* is indigenous to New Zealand

Our results confirmed the hypothesis that the indigenous *P. pulmonarius* var*. aotearoa* is genetically distinct from imported international strains, especially in the single copy genes RPB1, RPB2 and Tef. Phylogenetic analysis of ITS did not support the same conclusion. However, it is known from other genera in Agaricales that an ITS sequence similarity above 97-98.5% is not always conclusive to delineate species. For example, ITS was not useful in resolving species within the genera *Amanita* (Davison et al., 2017), *Armillaria* (Klopfenstein et al., 2017), *Pisolithus* and *Hygrocybe* (Badotti et al., 2017). Conversely, a higher threshold of >99% ITS sequence similarity was more conclusive than a 97% threshold for resolving species within the genus *Cortinarius* (Garnica et al., 2016). Our finding is further supported by a study on the larger *P. ostreatus* species complex using 40 single-copy genes, where the authors proposed that multiple species within *P. pulmonarius* exist, due to high intraspecific variation within those genes (Li et al., 2020). An earlier phylogenetic analysis using only ITS also showed several *P. pulmonarius* clades separated by country (Petersen & Hughes, 1997), supporting the discovery of a New Zealand indigenous *P. pulmonarius* strain a year before (Petersen & Ridley, 1996). The species delimitation metrics we tested were also employed in a study involving different clades of *Gymnopus confluens*, in which the authors applied naming of subspecies as a middle ground, in the absence of morphological taxonomic differences (Hughes & Petersen, 2015).

However, further research is required to fully ascertain whether *P. pulmonarius* var*. aotearoa* represents an endemic variety that is only found in New Zealand. There is a gap in knowledge about other Southern Hemisphere populations of this species. Therefore, we advocate for morphological taxonomic examination and DNA sequencing of new and existing *P. pulmonarius* collections from Australia, the Pacific Islands, South America, and other regions, amplifying, at minimum, the five gene regions employed in this study. Whole genome sequencing of a diverse range of *P. pulmonarius* specimens from around the world would further strengthen the evidence of speciation, possibly with a link to biogeography.

Whether or not *P. pulmonarius* var*. aotearoa* is also found in other countries, this clade represents collections from remote locations across New Zealand, indicating that multiple populations are well established here. Based on historical records of wild observations and the importation of international *P. pulmonarius* strains, it can be safely assumed that *P. pulmonarius* is indigenous to New Zealand rather than a recent arrival via anthropogenic dispersal. Our findings, which substantiate the claims of various New Zealand mushroom cultivators about the existence of an indigenous *P. pulmonarius* variety, may help increase the appeal and marketability of oyster mushrooms to New Zealanders by promoting the locally-grown, indigenous food aspect.

### P. purpureo-olivaceus has no anamorph and falls outside of Coremiopleurotus

The lack of evidence that historic or contemporary *P. purpureo-olivaceus* specimens produce asexual spores is supported by observations in a study on *Coremiopleurotus* that specifically investigated asexual reproduction (Zervakis, 1998). In the same study, Zervakis also suggested that *P. purpureo-olivaceus* differs from other species of *Coremiopleurotus* based on intercompatibility studies. Our finding that *P. purpureo-olivaceus* is not part of *Coremiopleurotus* (Figure 3) is also supported by a 25S rDNA phylogenetic study of Pleurotaceae (Thorn et al., 2000).

Why do two historic publications report an anamorphic stage of *P. purpureo-olivaceus*? The observation of coremia on a log without any *P. purpureo-olivaceus* sporocarps by Segedin et al. (1995) might relate to a different species and perhaps not a *Pleurotus.* Furthermore, the “minute spherical blackish structures” (Segedin et al., 1995) might have been spore-ball deposits from perithecial wood decay fungi, which are common on dead southern beech (McKenzie et al., 2000), or faeces of mites and other invertebrates. Only re-examination and ideally DNA sequencing would be able to confirm the claim by Segedin et al., but unfortunately, the two “anamorph” vouchers are lost. Petersen’s (1992) observation of “occasional ellipsoid arthrospores” in culture only occurred in one out of nine of his single-spore isolates of *P. purpureo-olivaceus* ICMP 9629. His observation indicates that if *P. purpureo-olivaceus* indeed produces asexual spores, their occurrence is rare. Given that we examined a much larger number of specimens than Petersen and Segedin et al. combined, the weight of the evidence indicates that *P. purpureo-olivaceus* does not have an anamorphic stage, unlike *P. australis*.

### Biosecurity implications of introduced strains of *P. djamor* and *P. pulmonarius*

The clear genetic differentiation between indigenous *P. parsonsiae* and exotic *P. djamor*, as well as between the two *P. pulmonarius* varieties, raises the question of whether import and cultivation regulations for *P. djamor* and exotic *P. pulmonarius sens. str.* in New Zealand should be revisited. Historically, the name *P. parsonsiae* was used as a synonym for *P. opuntiae* and *P. djamor* (Segedin & Pennycook, 2001; Zervakis et al., 2019), which is why *P. djamor* was deemed present in New Zealand prior to the legislation act in 1996 that restricted the importation of new organisms (Hazardous Substances and New Organisms Act 1996). Morphologically, genetically and ecologically, the true *P. djamor* is very distinct from *P. parsonsiae (Hosaka et al., 2024; Segedin et al., 1995; Stamets, 2000)*. To date, *P. djamor* has not been observed in the wild in New Zealand, possibly owing to its adaptation to tropical climates. *P. djamor* has a low tolerance for cold temperatures. We observed the senescence of mycelial cultures when refrigerated at 4 °C for several weeks.

However, as mean temperatures in New Zealand rise due to global warming, the risk of invasion of *P. djamor* into natural environments may increase. For example, a 2018 study in the United States showed how easily another cultivated oyster mushroom, *P. citrinopileatus* Singer, can spread and establish in the wild (Bruce, 2018). This species has now established all over the US (Jesionka, 2023; A. Pringle and M. Jusino, pers. comm.), with a study under way at University of Wisconsin on the impact of the spread of *P. citrinopileatus* on other wood-decay fungi and forest ecosystems linked to standing dead wood (Veerabahu et al., 2024). Rapid spread was also observed for *Favolaschia claudopus*, an invasive wood decay mushroom first discovered in New Zealand in 1969 and which has since spread throughout the country (Johnston et al., 2006). *F. claudopus* reproduces homothallically, a strategy that was linked to the invasion success of *Amanita phalloides* in the US (Wang et al., 2023). Nonetheless, the invasion risk of *F. claudopus* was initially suggested to be only moderate as it had been shown to be weakly competitive against indigenous saprotrophic fungi, instead thriving mostly in forest remnants and disturbed habitats where indigenous species may be less dominant (Johnston et al., 2006; Vizzini et al., 2009). This no longer appears to be the case because *F. claudopus* has become widespread in indigenous forests (N. Siegel, pers. comm.; https://www.inaturalist.nz, accessed 16/07/2024).

To reduce the risk that *P. djamor* may pose to New Zealand’s biodiversity, policymakers could consider the taxonomic complexity and invasion risks posed by *P. djamor*. It would be helpful for citizen scientists to report any wild observations of *P. djamor* to iNaturalist, MPI, or the authors of this study. The invasion risk will likely be driven by spore dispersal from small-scale home and commercial cultivation, which allows for genetic recombination, mutations and hybridisation with closely related indigenous species. Conversely, vegetative mycelial growth from discarded *P. djamor* growth blocks is less likely to lead to successful invasion. More research is needed to confirm this, as invasion strategies can vary between different fungi (Gladieux et al., 2015).

Our limited dataset on wild *P. pulmonarius* collections indicates that none of the international strains have escaped cultivation, although the absence of evidence is not conclusive evidence of absence. Exotic and indigenous *P. pulmonarius* are known to be sexually compatible (Petersen & Ridley, 1996); therefore, it is plausible that exotic or hybrid *P. pulmonarius* populations have established in the wild since this species was introduced for commercial and home cultivation in the 1990s. More collections and multi-gene sequencing would be required to map the potential spread of exotic *P. pulmonarius* strains.

## Conclusion

This study illustrates how systematics can impact biosecurity decisions designed to protect indigenous ecosystems. Using existing and new collections of *Pleurotus*, we established an updated phylogeny of this genus in New Zealand. We showed that a multi-locus analysis using ITS-LSU-RPB1-RPB2-Tef was necessary to confidently delineate species within Pleurotaceae, demonstrating the limitations of using ITS alone. Our results established that *P. pulmonarius* in the wild is indeed indigenous to New Zealand and separate at the level of variety from the introduced *P. pulmonarius* sens. str.; we proposed *P. pulmonarius* var*. aotearoa* as a new variety. Furthermore, we confirmed that *P. purpureo-olivaceus* falls outside the subgenus *Coremiopleurotus* and exhibits no anamorphic stage. The close sister relationship between *P. parsonsiae* and *P. djamor* underscores the need to reconsider the presence of the exotic *P. djamor* in the country. In summary, our study showed that the four indigenous *Pleurotus* species of New Zealand are distinct from each other and from overseas species. Our findings may help leverage more sustainable and competitive indigenous oyster mushroom cultivation practices in New Zealand.

## Supporting information

Supplement C - Expanded Methods

## Acknowledgements

We thank Carina Davis, Ana Podolyan, Katherine Trought, Dukchul Park, Diana Lee, Rose Williams, Amy Vaughan, Marion Donald, Panimalar Vijayan, Chris Winefield, Craig Herbold, and others in the School of Biological Sciences at University of Canterbury and at Manaaki Whenua for support, technical and otherwise. Anne Pringle and David Orlovich gave invaluable feedback on the thesis that led to this paper. We also thank the Department of Conservation for permission to sample (permit number CA-31615-OTH), and their staff for facilitating collections. This research was funded by University of Canterbury’s Food Transitions 2050 PhD scholarship, with additional funding from Ministry of Primary Industries through the MPI Postgraduate Science Scholarship 2023.

## Author Contributions

Funding was obtained by IAD, MKD, and PKB, with additional funding obtained by DH. All authors contributed to the study conception and design. Material preparation, data collection and analysis were performed by DH. Taxonomic examination and description was performed by JAC. The first draft of the manuscript was written by DH and all authors commented on previous versions of the manuscript. All authors read and approved the final manuscript. IAD, MKD and PKB were supervisors of DH’s PhD project.

---

Figure 5: Distribution map of Pleurotus records in Aotearoa. Data sources: New Zealand Fungarium, Te Kohinga Hekaheka o Aotearoa (PDD), International Collection of Microorganisms from Plants (ICMP), and iNaturalist (https://www.inaturalist.nz/). Sources are marked by filled diamonds (collections from the present study), squares (PDD), circles (ICMP), triangles (“Research Grade” iNaturalist observations), and open diamonds (genus level iNaturalist observations). Species are marked by colour: P. australis (brown), P. parsonsiae (tan), P. pulmonarius (blue) and P. purpureo-olivaceus (green). All records are dated: 16/07/2024

Figure 6: Multi-gene phylogenetic tree of New Zealand (NZ) Pleurotus, using maximum likelihood and Bayesian inference of concatenated ITS-LSU-RPB1-RPB2-Tef alignment. Numbers next to nodes denote bootstrap support/posterior probabilities. Taxa not found in NZ are marked in grey. Sequences from wild NZ collections are highlighted in yellow and imported taxa are highlighted in blue. Cultivated P. pulmonarius taxa isolated from the wild are marked with a star. Hohenbuehelia sequences were included to inform the position of P. purpureo-olivaceus within Pleurotus. The outgroup taxa (collapsed here for graphical representation) were Hygrocybe conica, Pterula gracilis, Sarcomyxa edulis and Typhula micans. Taxa in italics are missing sequencing data in one or more gene regions. Clades belonging to the subgenus Coremiopleurotus are marked with a vertical bar. All NZ P. purpureo-olivaceus strains are from the South Island, except those marked “North” for North Island collections.

Figure 7: P. pulmonarius var. aotearoa (Holotype, PDD 124119) micro morphological features in 5% KOH. (A) lamellar edge showing cheilocystidia; (B) basidia; (C) spores.

## Notes

### Competing Interest Statement

The authors have declared no competing interest.

