## Supplement B - Figures for "Multi-gene phylogeny and morphology of *Pleurotus* in Aotearoa New Zealand reveal a new variety of *Pleurotus pulmonarius*"

### Supplementary figures

| 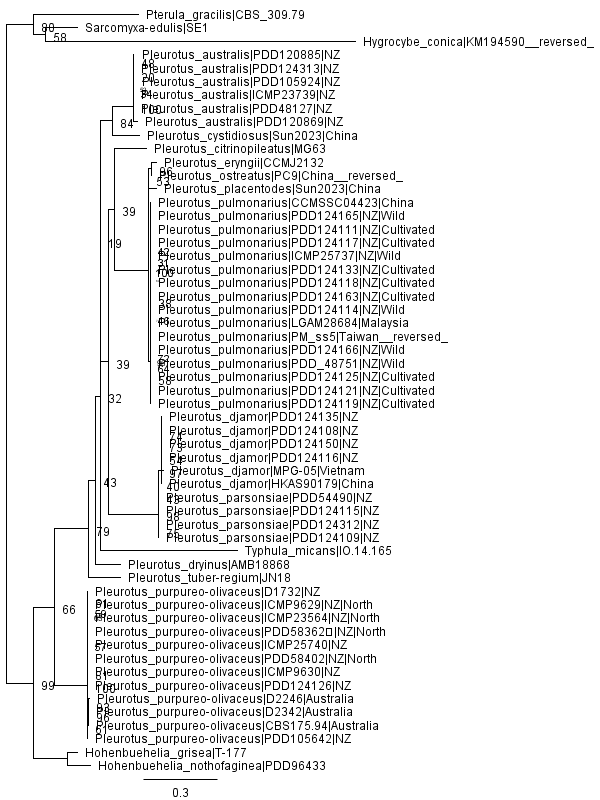 | 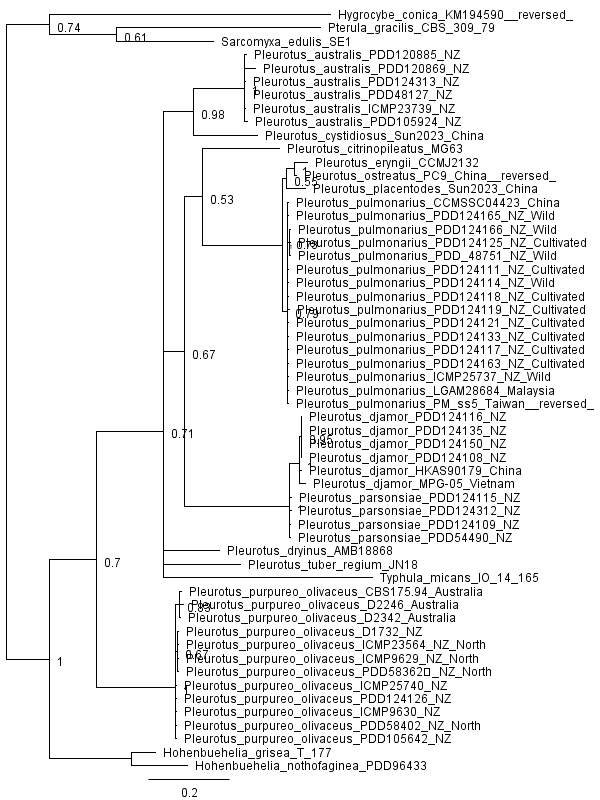 |
| --- | --- |

Figure B1: ITS phylogeny by ML (left) and Bayesian inference (right).

| 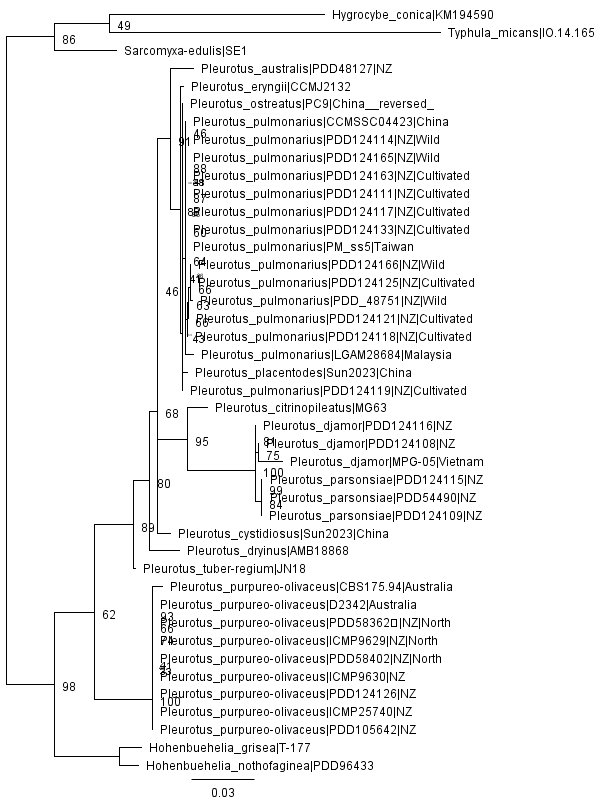 | 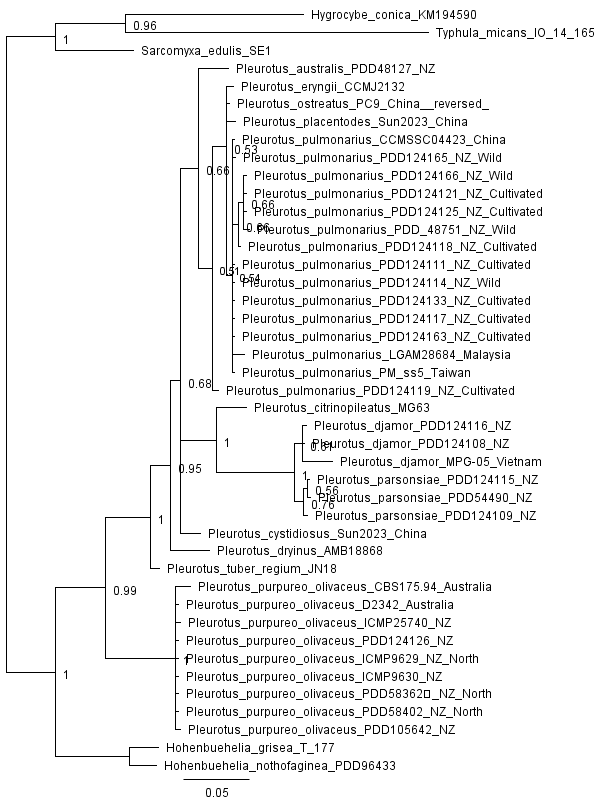 |
| --- | --- |

Figure B2: LSU phylogeny by ML (left) and Bayesian inference (right).

| 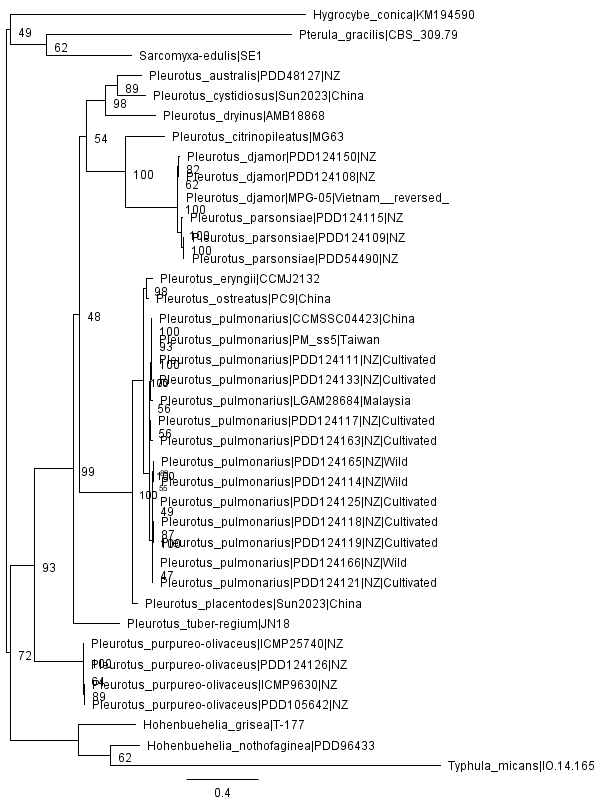 | 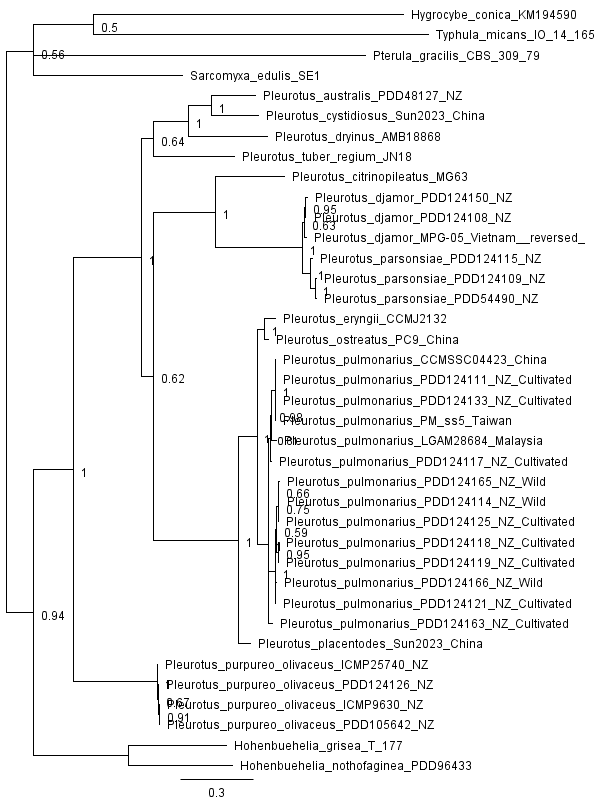 |
| --- | --- |

Figure B3: RPB1 phylogeny by ML (left) and Bayesian inference (right).

| 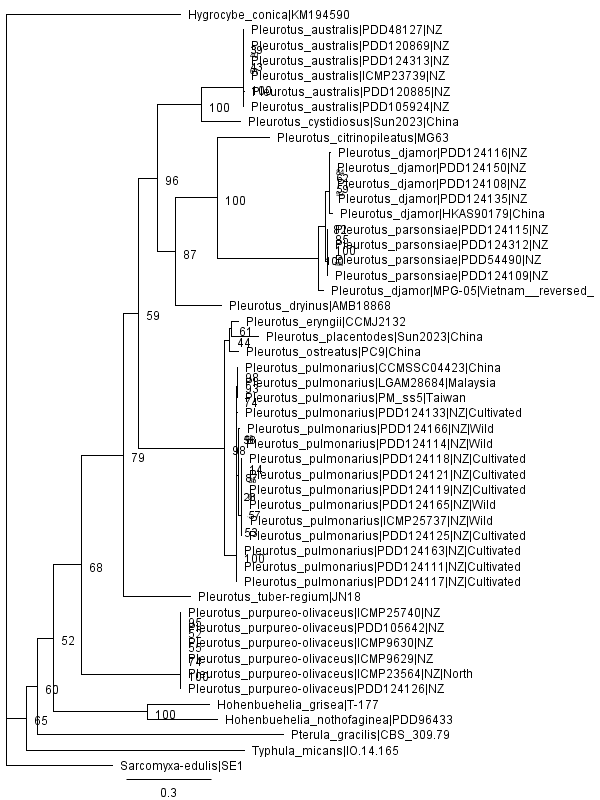 | 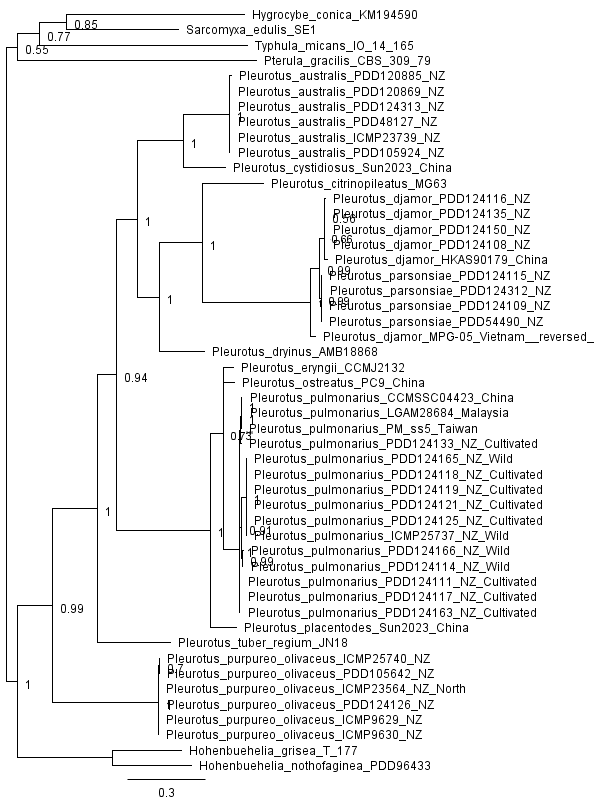 |
| --- | --- |

Figure B4: RPB2 phylogeny by ML (left) and Bayesian inference (right).

| 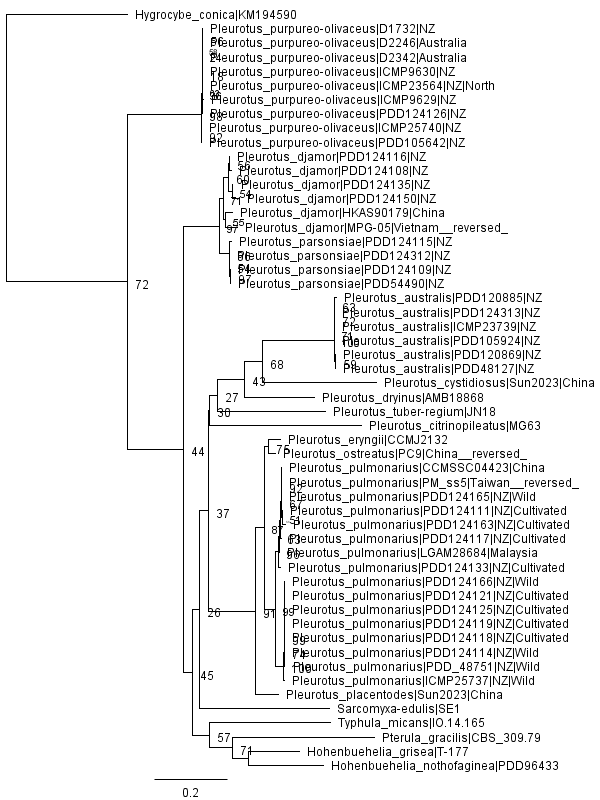 | 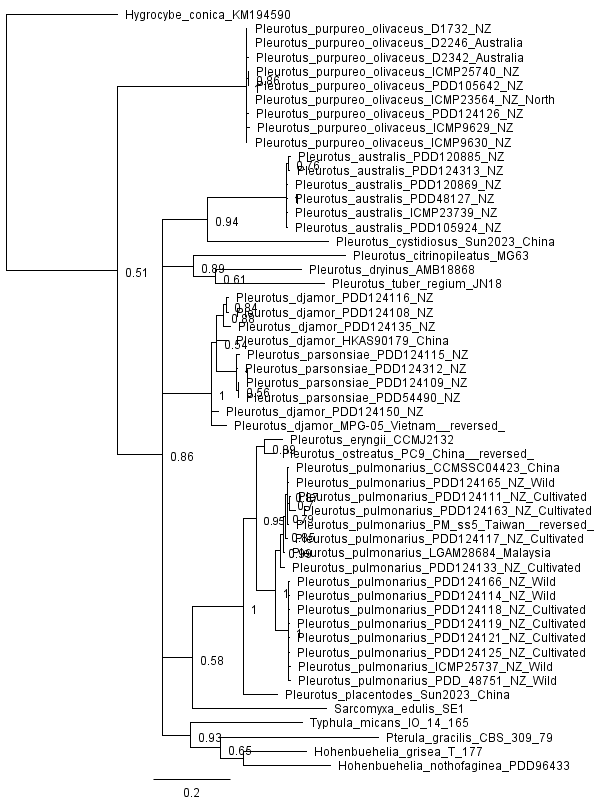 |
| --- | --- |

Figure B5: Tef phylogeny by ML (left) and Bayesian inference (right).
