## Supplement C - Expanded Methods for "Multi-gene phylogeny and morphology of *Pleurotus* in Aotearoa New Zealand reveal a new variety of *Pleurotus pulmonarius*"

### Extended Methods

#### Biological material

We collected 84 *Pleurotus* specimens from New Zealand for this study (Table A1). 43 specimens were sampled from the wild in 2021 and 2022, 19 strains were sourced from International Collection of Microorganisms from Plants (ICMP) and 22 strains were sourced from commercial suppliers of fresh oyster mushrooms, grow kits or inoculum. Collections for this study were permitted by the Manaaki Whenua - Landcare Research global concession from the Department of Conservation (number CA-31615-OTH) and with approval of the Ngāi Tahu Consultation and Engagement Group (as per their letter of May 11^th^ 2021). Additional collections in and around Auckland were covered by local Manaaki Whenua - Landcare Research permits from Auckland Council.

For sampling, we treated all sporocarps on the same individual host substrate as belonging to the same fungal strain. Each specimen comprised five intact sporocarps (or fewer if there were less than five present), ideally representing a variety of maturity stages and sizes, along with residual substrate at the base of the stipe. We recorded GPS coordinates, took *in situ* photographs before and after sampling, and logged observations on iNaturalist (<https://www.inaturalist.nz>). The majority of specimens were identified macroscopically, while some that looked similar to *Hohenbuehelia* spp. required microscopic examination. *Hohenbuhelia* is distinguished from *Pleurotus* by the presence of metuloid (thick walled) cystidia. The key macroscopic distinguishing traits of New Zealand *Pleurotus* species are cap colour, cap texture, gill colour, host substrate, stipe length, sporocarp growth pattern (solitary or fasciculate), and the presence of stipe and coremiospores (Segedin et al., 1995). Additionally, microscopic features such as spore shape, spore size and the presence of skeletal hyphae help distinguish macroscopically similar *P. parsonsiae* from *P. pulmonarius*, and to differentiate from other pleurotoid fungi.

We prepared cultures of field collections from sporocarp tissue to obtain dikaryotic cultures. We isolated tissue from the largest and least decayed sporocarp of each collection onto potato dextrose agar (Zervakis et al., 2019) supplemented with streptomycin to reduce contamination, followed by incubation at 25 °C in darkness. Pure cultures were stored as agar plugs in sterile water at 4 °C (Marx & Daniel, 1976). Samples in the form of mycelium from commercial suppliers were isolated as described above, and subsequently cultivated to produce sporocarps for storage as dried vouchers.

All material used in this study was deposited as voucher specimens at the New Zealand Fungarium, Te Kohinga Hekaheka o Aotearoa (PDD) in Auckland, and a subset in culture at ICMP in Auckland.

To investigate evidence of an anamorphic stage of *P. purpureo-olivaceus*, we macro- and microscopically examined the type collection and vouchers collected by Segedin et al. (1995). We further examined the culture ICMP 9629 described by Petersen (1992), as well as all specimens observed in the wild and in culture.

#### Taxonomic description

Dried material of *P. pulmonarius* specimens was soaked in 5% KOH before sectioning and microscopic examination employing Differential Interference Contrast (DIC) illumination. Measurements were taken from fruitbodies collected in the wild, and from cultivation of both wild-type and imported strains. Statistics derived from fruitbodies that are genetically identical were combined using the standard formulae for the mean of means. Standard colours are stated according to Kornerup & Wanscher (1967).

#### DNA extraction, PCR ampliﬁcation, sequencing and data assembly

We generated biomass for DNA extraction by growing cultures in custom liquid media (35 g l^-1^ glucose, 5 g l^-1^ peptone, 4 g l^-1^ yeast extract, 1 g l^-1^ KH_2_PO_4_, 0.5 g l^-1^ MgSO_4_) for 2 weeks, followed by washing with deionised water and squeezing out the material using sterile Miracloth (Merck Millipore, Germany). A manual cetyltrimethylammonium bromide (CTAB) based DNA extraction protocol (adapted from Jones & Schwessinger, 2021) was followed to generate high molecular weight DNA for Sanger sequencing and short-read whole genome sequencing. Briefly, fungal tissue was macerated in liquid nitrogen and lysed with CTAB buffer, RNAse A and Proteinase K (both from Macherey-Nagel, Germany) for 90 min at 60 °C in an orbital shaking incubator (Ratek OM11, Australia). As an additional step after lysis, samples were centrifuged for 5 min to precipitate proteins and other tissue that caused issues in DNA quality and yield during extraction from *Pleurotus* tissue. The supernatant was cleaned up twice with chloroform:isoamyl alcohol (24:1, v/v), followed by DNA precipitation with 100% ethanol and sodium acetate (3 M) at -20 °C overnight. Following centrifugation, the DNA pellets were washed twice with 75% ethanol, air-dried and eluted in Tris-HCl buffer (10 mM, pH 9.0). DNA quality was estimated using a NanoPhotometer NP80 (Implen GmbH, Germany), and DNA quantity was estimated using a Qubit 2.0 fluorometer (Invitrogen, USA) with the manufacturer’s broad range dsDNA BR assay kit. DNA extraction from nine samples failed to produce sufficient quality DNA for subsequent steps, dropping the total number of samples for sequencing to 75.

To investigate the phylogenetic relationships among *Pleurotus* samples, we analysed five molecular markers: the internal transcribed spacer region (“ITS” here, referring to ITS1+5.8S+ITS2) and the large subunit (LSU) of the nuclear ribosomal RNA gene, the first (RPB1) and second subunits of RNA polymerase II (RPB2), and the translation elongation factor alpha (Tef). The ITS region is often used for species identification of fungi but can be inconclusive for the phylogenetic distinction of closely related species in *Agaricales* (Davison et al., 2017; Klopfenstein et al., 2017). ITS sequences show significant degrees of intraspecific variability, which may lead to errors in species delimitation (Estensmo et al., 2021; Hilário et al., 2022; Nilsson et al., 2008; Tremble et al., 2020; Wilson et al., 2023). Two studies of *Pleurotus* found that Tef and RPB2 provided sufficient resolution to distinguish different species within the *Pleurotus eryngii* species complex (Estrada et al., 2010; Zhao et al., 2016).

From the DNA extracts, only ITS, RPB2 and Tef were amplified directly, whereas LSU and RPB1 were *in silico* extracted from whole genome assemblies as a complementary approach to add additional targets. PCR amplification was performed using primer pairs ITS4/ITS5 for ITS (White et al., 1990), fRPB2-5F/bRPB2-7.1R for RPB2 (Estrada et al., 2010), and TEF-983F/TEF-2218R for Tef (Suwannarach et al., 2020). PCR reactions were prepared with 1 µl DNA, 4 µl 5X Platinum II PCR Buffer, 0.4 µl dNTP (10 mM), 1 µl BSA, 0.8 µl of each primer (5 µM), 0.16 µl Platinum II *Taq* Hot-Start DNA Polymerase (ThermoFisher Scientific, USA) and 11.84 µl nuclease free water. The cycling conditions consisted of 2 min initial denaturation at 94 °C; followed by 35 cycles (ITS) or 40 cycles (RPB2 and Tef) of denaturation at 94 °C for 30 s; annealing at 52 °C (ITS) or 50 °C (RPB2 and Tef) for 30 s; extension at 68 °C for 30 s; 1 s final extension at 68 °C and 4 °C hold. Samples were run on electrophoresis gels with Lambda DNA/HindIII ladder (Thermo Fisher Scientific Inc., USA) to assess the size and successful amplification of each template. PCR products were sent to Ecogene (Auckland, New Zealand) for bidirectional Sanger sequencing. We auto-aligned and auto-trimmed reverse and forward sequences using Geneious 10.2.6 (Biomatters, Inc., New Zealand), with some low-quality sequences requiring additional manual editing and trimming using Sequencher 5.4.6 (Gene Codes Corporation, USA).

#### Library preparation, sequencing and *de novo* genome assembly

In addition to Sanger sequencing, we prepared 54 samples for short-read whole genome sequencing using the MGI platform (MGI Tech, China), in order to extract additional target gene region data for enhancing the phylogenetic analysis. We had to exclude 30 samples from genome sequencing because of either insufficient genomic DNA or due to limited consultation with local iwi (indigenous tribes). Two pooled libraries were constructed using the MGI Fast FS DNA Library Prep Set (MGI Tech, China), according to the manufacturer’s instructions. 51 strains were included in the first medium-coverage library, with one strain of each of *P. parsonsiae*, *P. pulmonarius* and *P. purpureo-olivaceus* used in the second ultra-high coverage library to facilitate *de novo* assembly of reference genomes. Briefly, 25-200 ng of DNA was enzymatically fragmented, end-repaired, adapter ligated, amplified using PCR (due to low input amount), purified, and quantified using Qubit. All PCR products were then pooled equimolar into a single library, and we used the MGIEasy Dual Barcode Circularization Kit (MGI Tech, China) for single strand circularisation and enzymatic digestion (as per manufacturer’s instructions), followed by making a DNA nanoball from 50 fmol of the constructed library. Subsequently, 150-bp paired-end sequencing of the DNA library was performed using DNBSEQ-G400 (MGI Tech, Beijing, China) on a single lane of an FCL-PE150 flow cell using a commercial sequencing service (Lincoln University, New Zealand).

Demultiplexed reads were quality trimmed to Q30 using BBMap v39.01 (Bushnell, 2023) and *de novo* assembled using SPAdes v4.0.0 (Bankevich et al., 2012) with default parameters. Assemblies of all *P. pulmonarius* and *P. djamor* strains were further improved with a reference guided assembly using RagTag v2.1.0 with default parameters of the scaffold function (Alonge et al., 2022) to *P. pulmonarius* reference genome GCA_012980535.1 (Vidal-Diez de Ulzurrun et al., 2021) and *P. djamor* reference genome GCA_029747585.1, respectively.

#### *In silico* target gene region extraction from genomes

Some key samples did not amplify well for ITS, RPB2 and/or Tef using Sanger sequencing with standard primers and PCR conditions; therefore, we used two different methods to *in silico* extract ITS, RPB2 and Tef from the *de novo* assembled genomes, with the addition of LSU and RPB1.

First, we utilised a BLASTn subject query (Camacho et al., 2009) to extract matches of representative regions of ITS, LSU, RPB1, RPB2, and Tef within each genome assembly using closely related reference sequences from GenBank for each species, where available. Second, we used ThermonucleotideBLAST v2.61 ("tntblast", Gans & Wolinsky, 2008) to extract predicted amplicons from each genome assembly using forward and reverse primers for each target region. Specifically, we used the primer pairs ITS5/ITS4 for ITS (White et al., 1990), LR05/LR5 for LSU (Zervakis et al., 2019), RPB1-Af/RPB1-Cr for RPB1 (Matheny, 2005), bRPB2-3.1F/gRPB2-6R and RPB2-5F/bRPB2-11R1 for RPB2 (Liu et al., 1999), and TEF-983F/TEF-1567R for Tef (Li et al., 2017). The BLASTn approach had a higher success rate but resulted in shorter amplicons than tntblast (Table B3).

To further strengthen the multi-gene phylogenetic analysis, we extracted the same loci from other *Pleurotus* reference genomes on NCBI, as well as several other closely related *Agaricales* taxa genomes (Table A2), in order to have full representation of the same loci across outgroups and ingroups.

Supplementary Table B3: Comparison of the three different methods used to extract target gene regions, using Sanger sequencing, in silico extraction using BLASTn target query, and in silico amplification with primer search using ThermonucleotideBLAST (“tntblast”). The mean length of extracted barcodes of n samples is given for each locus and method, along with the success rate (percentage of samples that resulted in a sequence that matched expected Pleurotus sequences) and the percentage of ambiguous base pairs for Sanger sequencing. Neither in silico approach resulted in ambiguous base pairs. LSU and RPB1 were only in silico extracted.

|  | Sanger sequencing | | | | *In silico*  BLASTn target query | | | *In silico*  primer search (tntblast) | | |
| --- | --- | --- | --- | --- | --- | --- | --- | --- | --- | --- |
| Gene region | n | Length | Success rate | Ambiguous base pairs | n | Length | Success rate | n | Length | Success rate |
| ITS | 75 | 598 | 83% | 0.03% | 50 | 555 | 100% | 50 | 652 | 80% |
| RPB2 | 75 | 1056 | 75% | 0.17% | 50 | 1130 | 96% | 50 | 2215 | 92% |
| Tef | 75 | 1040 | 68% | 0.11% | 50 | 1065 | 100% | 50 | 1204 | 100% |
| LSU |  |  |  |  | 50 | 832 | 100% | 50 | 974 | 58% |
| RPB1 |  |  |  |  | 50 | 366 | 100% | 50 | na | 0% |

#### Sequence alignment and phylogenetic analyses

We chose the *in silico* extracted amplicons for subsequent steps, supplementing them with amplicons from taxa which only Sanger sequencing data was available for. The genome-extracted amplicons were given preference due to the higher quality of reads and better representation of the target regions (Table B3), especially RPB2, for which only partial gene amplification was successful with Sanger sequencing using a single primer pair.

In addition to the sequences generated from the biological material in this study, sequences from ICMP and PDD were included, along with ITS, LSU, RPB1, RPB2 and Tef sequences of other taxa from GenBank. No international *Pleurotus* sequences in GenBank that were related to the New Zealand species had sufficient representation for all five gene regions.

Sequences were aligned separately for each region using MAFFT v7.450 and the E-INS-i algorithm (Katoh & Standley, 2013). Alignments were manually realigned in Geneious, to correct misaligned gaps across related taxa. A multi-gene alignment was generated by concatenating the five alignments using the “Concatenate Sequences or Alignments” function in Geneious.

Phylogenetic analysis was based on maximum likelihood and Bayesian inference for each individual gene region and the concatenated alignment. Maximum likelihood (ML) analyses were conducted with IQ-TREE v2.3.6 (Nguyen et al., 2015), using the -MFP command to automatically choose the appropriate substitution model for each partition (using the -p command). Substitution models were restricted to those compatible with MrBayes for the subsequent Bayesian analysis, using -mset mrbayes. Bootstrap values were computed using Ultrafast analysis of 1000 pseudoreplicates, to evaluate branch confidence and find the best-scoring ML tree.

Bayesian phylogenies were constructed using MrBayes v3.2.7a (Ronquist et al., 2012), using the optimal substitution models as identified by IQ-TREE ModelFinder for each partition (GTR+I+G for all partitions except for ITS2 and the first Tef intron) and converted to the appropriate MrBayes block configuration settings using PhyloSuite v1.2.3 (Xiang et al., 2023). Posterior probabilities were estimated using two parallel searches on four chains sampled every 1000 generations, run from a random starting tree for four million Markov chain Monte Carlo generations, to achieve an average deviation of split frequencies below 0.01. Branch lengths were linked across the two partitions. A burn-in setting was used to discard the first 25% of trees to construct a majority rule consensus tree using the “halfcompat” setting for “contype”.
